# Cold Comfort: metabolic rate and tolerance to low temperatures predict latitudinal distribution in ants

**DOI:** 10.1101/2023.02.11.527843

**Authors:** Quentin Willot, Michael Ørsted, Hans Malte, Johannes Overgaard

**Affiliations:** Department of Biology, Aarhus University, 8000 Aarhus C, Denmark; Department of Chemistry and Bioscience, Aalborg University, 9220 Aalborg E, Denmark

## Abstract

Metabolic compensation has been proposed as a mean for ectotherms to cope with colder climates. For example, under the metabolic cold adaptation/metabolic homeostasis hypotheses (MCA/MHH), it has been formulated that cold-adapted ectotherms should display higher/more thermally sensitive metabolic rates (MRs) at lower temperatures. However, whether such compensation can truly be associated with distribution, and whether it interplays with cold-tolerance to support species’ climatic niches, remains largely unclear despite broad ecological implications thereof. Here, we teased apart the relationship between MRs, cold-tolerance, and distribution, to confront the MCA/MHH among 13 ant species. We report clear metabolic compensation effects, consistent with the MCA and MHH, where MR parameters strongly correlated with latitude and climatic factors across species’ distributions. The combination of both cold-tolerance and MR further upheld the best predictions of species’ climatic niches. To our knowledge, this is the first study showing that the association of metabolic data with cold-tolerance supports increased predictive value for biogeographical models in social insects. These results also highlight that adaptation to higher latitudes in ants involved adjustments of both cold-tolerance and MRs, potentially at the expense of metabolic performance at warmer temperatures, to allow this extremely successful group of insects to thrive under colder climates.

## Introduction

Colder climates that usually define high-altitude/latitude environments pose two important challenges to cold blooded organisms. First, they entail the occurrence of cold-extremes and second, they constrain the amount of energy available for development and reproduction through higher seasonality [1]. Consequently, two commonly observed strategies observed in colder-adapted ectotherms are increased cold-tolerance and comparatively shortened seasonal activity cycles [2, 3], which has been argued to sometimes accompany metabolic adaptations [4-7]. Several hypotheses about ectotherms’ metabolic strategies to cope with cold climates have been formulated (Fig. 1), among which the metabolic cold adaptation hypothesis (MCA) [8] has probably met the most scrutiny. The MCA states that to exploit shorter annual timeframes for development, ectotherms inhabiting cooler environments should display higher standard metabolic rates (SMRs) to compensate for the effects of both lower temperatures and shorter growing seasons [4, 9, 10]. Another somewhat less assessed hypothesis layering on the MCA is the ‘metabolic homeostasis’ hypothesis (MHH) [11]. This directly relates to the thermal sensitivity of the SMR, often assessed by measuring Q_10-SMR_ values (the factorial change in SMR due to an increase of temperature by 10°C) over several temperature intervals. The MHH suggests that that cold-adapted ectotherms should also display high SMR sensitivity at a lower temperature range to quickly take advantage of mild warming episodes and, conversely, that they would also reach peak metabolic rates at lower temperatures than in their warm-adapted relatives [7, 12]. Both these hypotheses have been tested on a wide range of ectothermic organisms, but have remained, to this day, controversial mainly due to the lack of consistent support across taxa [4, 6, 7, 12-16].

**Figure 1.**
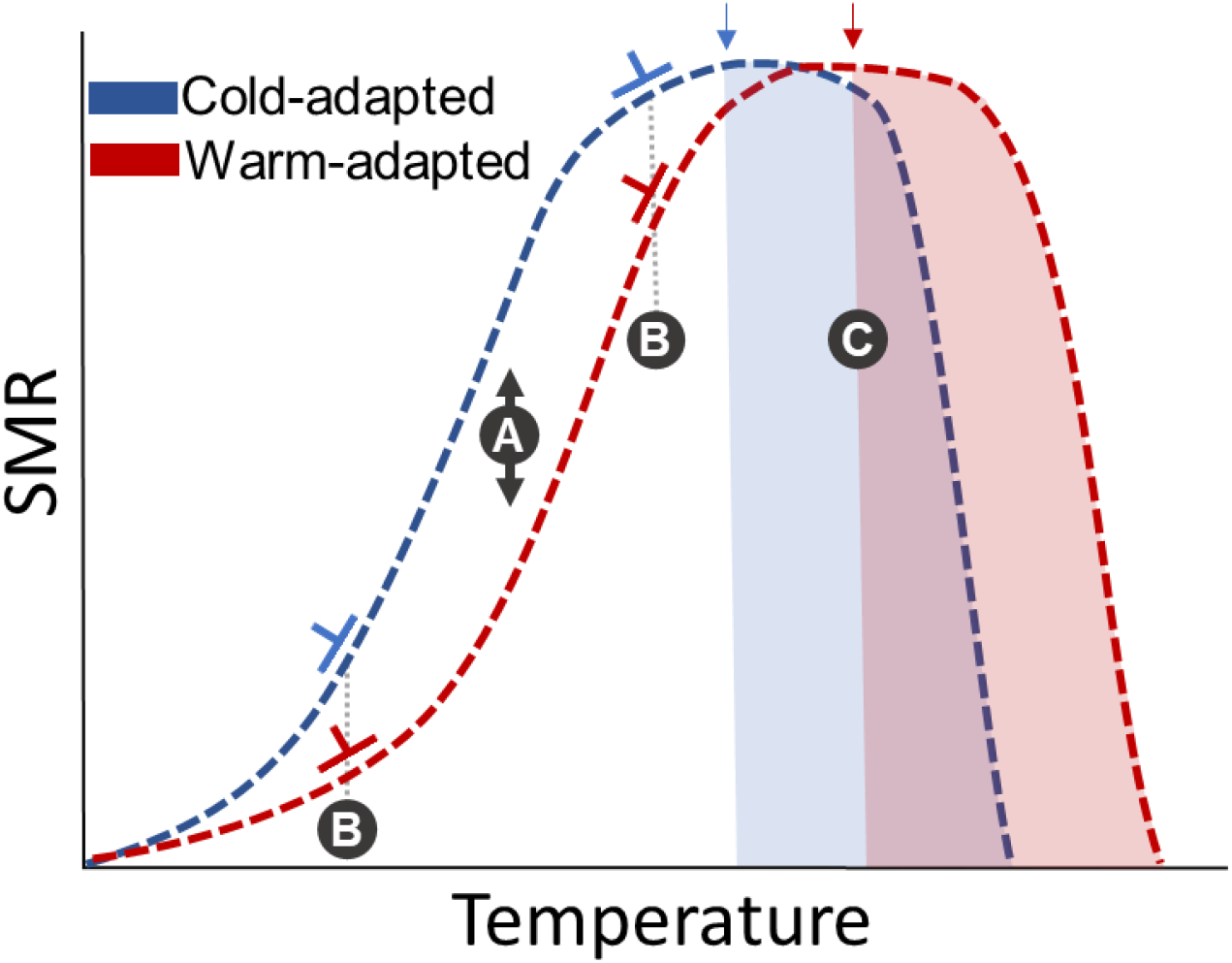
Hypothetical relationship between standard metabolic rate (SMR) and temperature for cold adapted (blue) and warm adapted species (red) of ectotherms under a metabolic compensation framework. **A**. The MCA hypothesis states that cold-adapted ectotherms should display higher SMR at lower ranges of temperature. **B**. The MHH states that the thermal sensitivity of the metabolic rate, related to the slope of this relationship (Q_10-SMR,_ represented by orthogonal lines), should be comparatively higher at lower temperature for cold adapted ectotherms. Conversely, cold-adapted ectotherms should also approach peak metabolic rates at lower temperatures than in their warm-adapted relatives, where they will display comparatively reduced Q_10-SMR_ values. **C**. Range of temperatures where gradual decline in fitness for cold (blue) and warm (red) adapted species might occur.

While the occurrence of metabolic compensation at larger scales in ectotherms is thus debated, increased cold-tolerance on the other hand has been highlighted as a hallmark of adaptation to colder-climate. The association between cold-tolerance and species’ distributions has found ample support in the literature, including in insects [17-21]. Interestingly, a putative functional link between cold-tolerance and metabolic compensation has been proposed [22], since increased MRs could also provide the energetic means for species to defend ion-balance and delay the onset of, as well as help recovering from, chill-induced coma [23]. Albeit still in need of substantial testing, this hypothesis provides an interesting framework that could bridge metabolic compensation, cold-tolerance, and distribution in ectotherms. It is still unclear whether metabolic compensation supports and/or interacts with cold-tolerance to drive ectotherms’ distribution, and whether metabolic compensation represents a taxa-specific strategy or a more general rule within the adaptative potential of ectothermic species. Addressing this fundamental but complex question has been at the heart of the exploration of ectotherm adaptation to colder climates for decades [5, 24, 25]. It might have become even more relevant within the context of climate change. Evidence taken from invertebrate species appear to place optimal temperature for traits such as growth slightly below temperatures for peak SMRs [26]. This suggests that displacements of SMR thermal performance curves (TPC) towards cooler ranges of temperatures through metabolic compensation could also in ectotherms trade off with the performance of other life-history traits at warmer ranges (Fig. 1, see [26] for review). In this study, we aimed to: (1) reestablish the well-known association between cold tolerance and latitudinal/climatic distribution in insect models, (2) test the occurrence of MCA/MHH within a single consistent phylogenetical framework of 13 ant species covering a wide latitudinal distribution in western Europe, (3) explore the climatic factors associated with metabolic compensation and cold tolerance using current biogeographical tools, and finally (4) test the potential implication of metabolic compensation in predicting species distributions and climatic niches, and whether we could characterize peak SMR temperatures in cold-adapted species. Ants represent an opportune model to undertake such exploration. Firstly, they initially radiated from their center of diversity in the tropics to colonize temperate and boreal climates [27] and have come to colonize the vast majority of terrestrial microhabitats [28], thus offering an relevant phylogenetic framework to tease apart the gradual evolution to colder climates in insects. Secondly, adults are comparatively long-lived and sessile [29, 30], thus having to endure their local seasonal climatic conditions sometimes over several years. Here, briefly, we report here a clear metabolic compensation effect in higher latitude species after phylogenetic correction of data in line with the MCA/MHH. We also show that the association of cold tolerance and metabolic rate, although not directly correlated, is best at predicting both latitudinal limit of distribution and mean environmental temperature over species’ distribution ranges. Finally, we show that Q_10-SMR (25-30°C)_ values for species were inversely correlated with latitudes in our model system, suggesting that colder-adapted ant might be much closer to peak-SMR in the 25-30°C temperature range than their Mediterranean relatives. This interval could thus represent an important tipping point in some temperate ant species, where they could be subjected to a gradual decline in fitness. Overall, this work shows that adaptation to a colder climate is a complex trait that in ants leveraged both shifts in cold-tolerance and SMR, allowing them to colonize and thrive in colder climates.

## Methods

### Animal model system, field sampling, and laboratory rearing

Species sampling and laboratory rearing were performed as described previously [31]. Briefly, thirteen ant species from common genera in western Europe were collected to represent a diversity of latitudinal distribution. Colonies of *Lasius niger, Lasius flavus, Formica fusca, Myrmica rubra*, and *Leptothorax acervorum* were collected near Aarhus, Denmark (56.16°N, 10.20°E), which is characterized by a hemiboreal climate (Köppen classification: Dfb). Colonies of all other species (*Lasius cinereus, Lasius grandis, Lasius emarginatus, Lasius myops, Aphaenogaster senilis, Aphaenogaster gibbosa, Aphaenogaster subterranea, Cataglyphis piliscapa*) were collected around Collioure, France (42.52°N, 3.08°E), which is characterized by a Mediterranean climate (Köppen classification: Csa). Colonies were kept in plastic boxes with Fluon®-coated sides and provided with16×150 mm plastic test tubes with a water-filled section in the bottom separated with a moist cotton plug for nesting. All colonies were kept for at least two months prior to experimentation under a 12:12 light:dark cycle at constant 26°C, and fed honey water and sliced mealworms twice a week.

### Dynamic cold tolerance assays

Cold knockdown temperature was tracked by recording critical thermal minimum (CT_min_) on workers placed individually into 5 ml closed glass vials. Glass vials were mounted to a rack and submerged into a transparent tank filled with a 25% ethylene-glycol:water solution with a programmable water bath (LAUDA-Brinkmann, NJ, USA). Temperature was maintained at 20°C for 15min to allow workers to acclimate, and then gradually decreased at a rate of 0.1°C/min until knockdown that was scored by a total absence of movement of workers, even after external stimulation (gentle vial shaking). CT_min_ was recorded as the average knockdown temperature for each species (N=10-20 workers per species).

### Metabolic rates measurements

Metabolic rates of our 13 ant species were estimated indirectly from the rate of CO_2_ production (VCO_2_) using repeated measurements of stop-flow respirometry [32, 33]. Here, we used an experimental setup similar to that described by Jensen et al., 2014 [34]. Twenty workers of each species were randomly assigned to one of 6-parallel cylindrical glass chambers (20 mm diameter, 70 mm length). Within each chamber, workers had access to a 1.5ml Eppendorf tube filled with 10% honey water to prevent starvation or desiccation during measurements. For each experimental run, respiration chambers were kept in the dark inside a programmable temperature cabinet (Termaks cabinets; KB8400m, GmbH, Austria). The temperature inside the cabinet was programmed to cycle by 8 hours periods, from an initial 25°C to 18°C and then to 30°C over a 24h experimental period. Thus, VCO_2_ of workers was recorded at each temperature for approx. 8h (See Fig. S1). Two parallel 8-channel-multiplexers (RM Gas Flow Multiplexer, Sable Systems, USA) controlled the sequential flushing and closing of the respiration chambers such that the stop-flow respirometry system obtained repeated measures of VCO_2_ in the 16 parallel respiration chambers (typically 12 chambers with ants and 4 empty chambers). Airflow was controlled by a mass flow meter (Side-Trak, Sierra Instruments, USA), regulated by a flow controller (MFC 2-channel v. 1.0, Sable Systems, USA). During the flush phase (180 s) the respiration chambers were perfused with CO_2_ stripped air (air passing a soda lime column (MERCK Millipore, Darmstadt, Germany) at a fixed rate of 150 mL*min^-1^. After the flush phase the respiration chamber was closed for 2700 s to allow CO_2_ to accumulate, while the remaining 15 chambers were flushed sequentially in a similar manner. Accordingly, a CO_2_ peak accumulated during the closed time was measured from each chamber every 2880s (ca. 48 min). Air leaving the respiration chambers passed a calcium chloride column (AppliChem, Darmstadt, Germany) to remove moisture before entering a CO_2_ analyzer (Li700 CO_2_ Analyzer, LI-COR Environmental, Lincoln, Nebraska, USA). Data were sampled at 1-s intervals using an UI2 Sable Systems interface and collected using ExpeData software (ExpeData-UI2 software, Sable Systems, Las Vegas, Nevada, USA). VCO_2_ values were obtained by the automated integration of the area beneath each peak using a script in Mathematica 7 (Wolfram Research, Champaign, Illinois, USA) [34].

On completion of measurements, the total fresh weight of workers was measured to the nearest 0.1 mg (Sartorius, Type 1712, Göttingen, Germany) to allow for calculation of mass specific metabolic rate. The first two rounds of measurement as well as measurements during temperature transitions were discarded to allow workers to settle down in the respiration chamber (ca. 90min to transition between temperatures, Fig. S1). Finally, chambers displaying more than a 10% mortality rates (2 workers) following the 24h run were excluded form subsequent analysis. Following these steps, we selected the lowest 3 values at each temperature for each chamber, counting 3 to 6 chambers per species. (Fig. S1).Two chambers were kept empty and CO_2_ accumulation in these chambers resulting from unwanted diffusion (typically less that 2-3% of chamber with ants) was subtracted from that measured in chambers with ants.

To estimate resting/standard MR, VCO_2_ values were corrected for the fresh mass of ants involved in the experiments, and we adjusted for size allometry between species using a scaling exponent of 0.75 [35]. In *Drosophila*, it was demonstrated that this method of repeated stop flow respirometry produces MR estimates that are generally slightly lower than measurements obtained using open flow protocols [16, 34]. Finally, we used the equation Q_10_=(R_2_/R_1_)^10/(T2-T1)^ to estimate Q_10-SMR_ values, where R_2_ and R_1_ are the respiration rates at temperature T1 and T2, respectively [14]. We calculated Q_10-SMR_ values at two temperature ranges: 18-25 and 25-30 °C.

### Phylogenetic signal testing

We tested phylogenetic signal as described previously [31]. Briefly, we used the genus-level ant phylogeny [36] pruned to genera level and then compiled a time-calibrated phylogeny of *Lasius [37]* to reconstruct the phylogeny of *L. niger, L. emarginatus, L. flavus, L. myops after which we* manually added *Lasius cinereus* and *Lasius grandis* at their right relative position within the genus [38]. Similarly, we placed Aphaenogaster species at their right relative position based on Schifani *et al*., 2022 [39]. We then used Pagel’s λ [40] to test for phylogenetic signal in CT_min_, and SMR at our three tested temperatures using the packages ape [41] and phytools [42].

### Biogeography

We tested for the putative association between cold tolerance, mass-corrected SMR, Q_10_-_SMR_, and climatic variables using climatic variables averaged over the species full distribution ranges as described in Willot *et al*., 2022 [31]. Briefly, species’ occurrence data were extracted from the Global Ant Biodiversity (GABI) database [43]. Only georeferenced records were used and were restricted from longitude 30°W to 145°E and latitude 30°N to 72°N. We matched soil temperature bioclimatic variables from recent maps of global soil temperature at 0-5cm depth [44], and precipitation variables from the WorldClim v2.1 database [45] with species’ distributions. All bioclimatic variables were used in a 30 arc second resolution (approx. 1×1 km at the equator). These maps were projected to a Behrmann’s cylindrical equal-area projection with true scale along latitude 50°N. To account for sampling bias, we aggregated these values to a 20-arc min resolution (approx. 40×40km), *i*.*e*., all 1×1km observations within the same cell were averaged. We then used a Brownian motion co-variance linear model accounting for phylogenetic signal using the R-package Caper (Orme *et al*., 2018) to regress values of species CT_min_, SMR and Q_10-SMR_ against soil and precipitation climatic variables.

### Data analysis

Phylogenetic signal testing, biogeographical correlations using Phylogenetic Generalized Least Squares models, Principal Component Analysis and model testing through Akaike Information Criterion (AIC) analysis were performed in R version 4.1.2 [46].

## Results

### Cold tolerance tracks environmental climate and distribution

CT_min_ values (mean ± SD) ranged from -4.14±0.47°C (*L. acervorum*) to 1.09±0.49°C (*A. subterranea*; Fig. 2A). There was a moderate but not statistically significant phylogenetic signal for CT_min_ (λ=0.93, *p*=0.140) that was taken into account for all subsequent correlations involving cold-tolerance. CT_min_ correlated strongly with northern extremes of latitudinal distribution and with several soil (for temperatures) and air (for precipitations) climatic variables averaged over species’ distributions (Fig. 2B, C, Table 1). This indicates that species living under on average colder temperatures and higher latitudes had lower CT_min_, and increased cold-tolerance. The best predictors of species CT_min_ were annual mean soil-surface temperature (r=0.873; *p*<0.001, Fig 2.B) and northernmost limit of latitudinal distribution (r=-0.816; *p*<0.001 Fig 2. C).

**Fig. 2.**
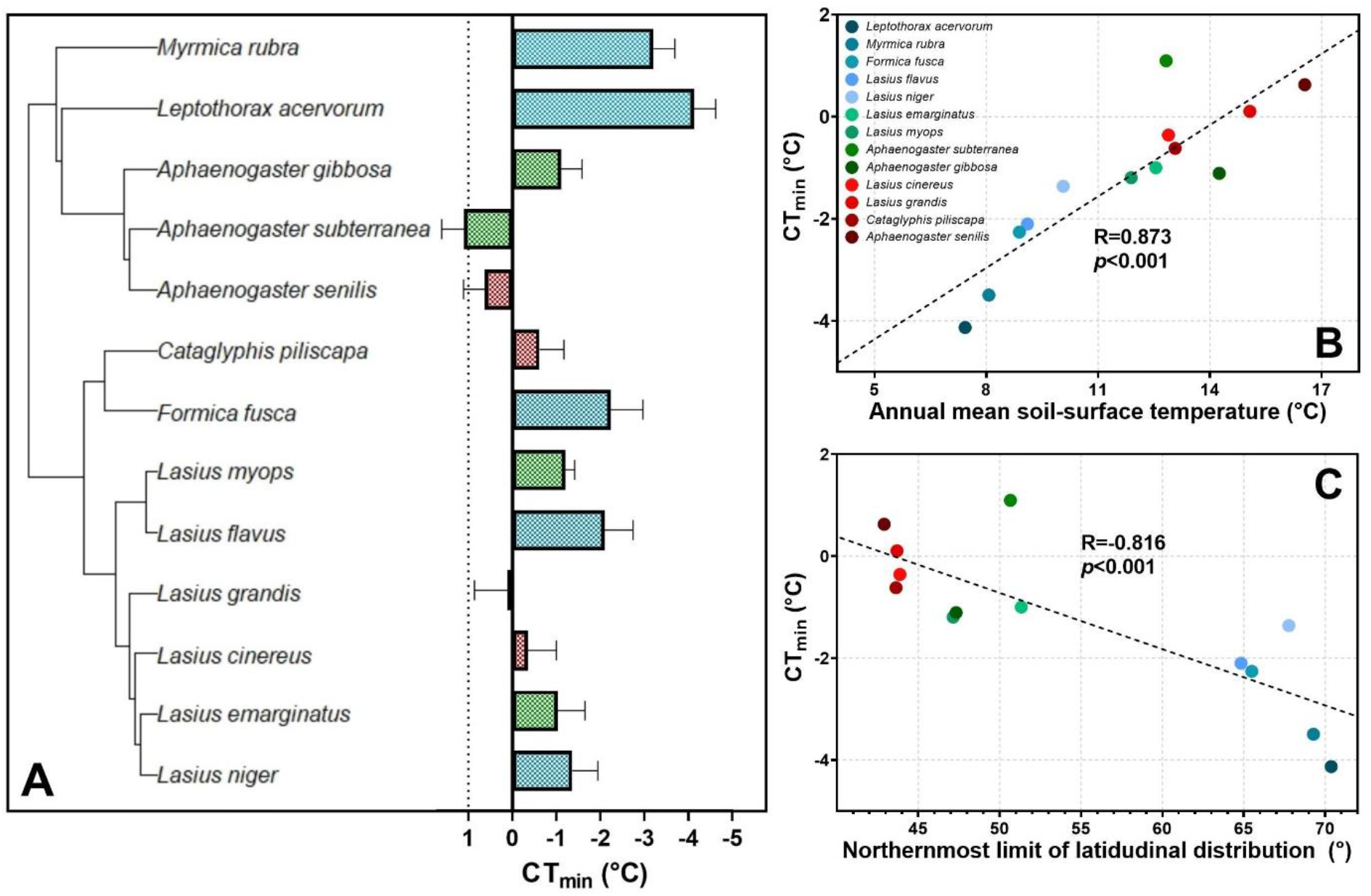
Cold-tolerance tracks annual mean soil-surface temperatures and northernmost limits of latitudinal distribution. Species are color coded as being either Mediterranean (red) or extending their range towards intermediate (green) or northward (blue) latitudes within continental Europe. **A**. CT_min_ of species mapped across the phylogeny of our model system. **B**. Phylogenetically-corrected correlation between species CT_min_ and annual mean soil-surface temperatures (r=0.873, *p*<0.001). **C**. Phylogenetically-corrected correlation between species CT_min_ and northernmost limit of latitudinal distribution (r=-0.816, *p*<0.001).

**Table 1.** Phylogenetically corrected correlations between CT_min_ of the 13 ant species included in our analysis regressed against extremes of latitudinal distribution and several climate variables averaged over species’ distributions. Temperature variables represent soil temperature at a 0-5cm depth [44] and precipitation variables are drawn from the WordClim database [45]. The strength of these correlations is presented in shades of red (Pearson’s correlation coefficient, r values), and its statistical significance in shades of blue (*p* values). Overall, there was a very strong correlation between species CT_min_ and annual mean soil-surface temperature (r=0.873; *p*<0.001, Fig 2.B) as well as northernmost limit of latitudinal distribution (r=-0.816; *p*<0.001 Fig 2. C)

| Climatic variables | CT<sub>min</sub> |  |
| --- | --- | --- |
|  | r | p |
| Annual mean soil temperature | 0.873 | <1*10<sup>-3</sup> |
| Northernmost latitude of | -0.816 | <1*10<sup>-3</sup> |
| Min soil temperature | 0.794 | <1*10<sup>-2</sup> |
| Max soil temperature | 0.780 | <1*10<sup>-2</sup> |
| Annual precipitation | -0.742 | <1*10<sup>-2</sup> |
| Precipitation seasonality | 0.673 | 0.011 |
| Mean soil temperature diurnal | 0.573 | 0.040 |
| Southernmost latitude of | 0.152 | 0.620 |

### Metabolic rate at lower temperatures tracks distribution

Mass-corrected SMRs were assayed at 18, 25 and 30°C for the 13 species included in our analysis (Fig. 3A, Fig. S.2). Most species extending their range northward in colder and rainier habitats exhibited on average significantly higher SMRs at 18°C than Mediterranean species (Fig. 3A). This trend remained rather consistent at 25°C but disappeared at 30° (Fig. 3A). We found moderate but not statistically significant phylogenetic signals in SMRs for respiration rates at trial temperatures, nevertheless these were accounted and corrected for in subsequent correlations. Overall, northernmost limits of distribution as well precipitation and temperature variables were good predictors of SMRs at lower but not higher temperatures (Table 2). The best predictors of species mass-corrected SMRs at 18°C were northernmost limits of latitudinal distribution (r=0.674, *p*=0.011, Fig. 3B) and annual precipitation (r=-0.665, *p*=0.013, Fig. 3C). Consistently, these predictors lost power but remained statistically significant at 25°C, while none of them were significant at 30°C (Table 2). In all cases, phylogenetically corrected correlations between CT_min_ and SMRs yielded poor associations.

**Fig. 3.**
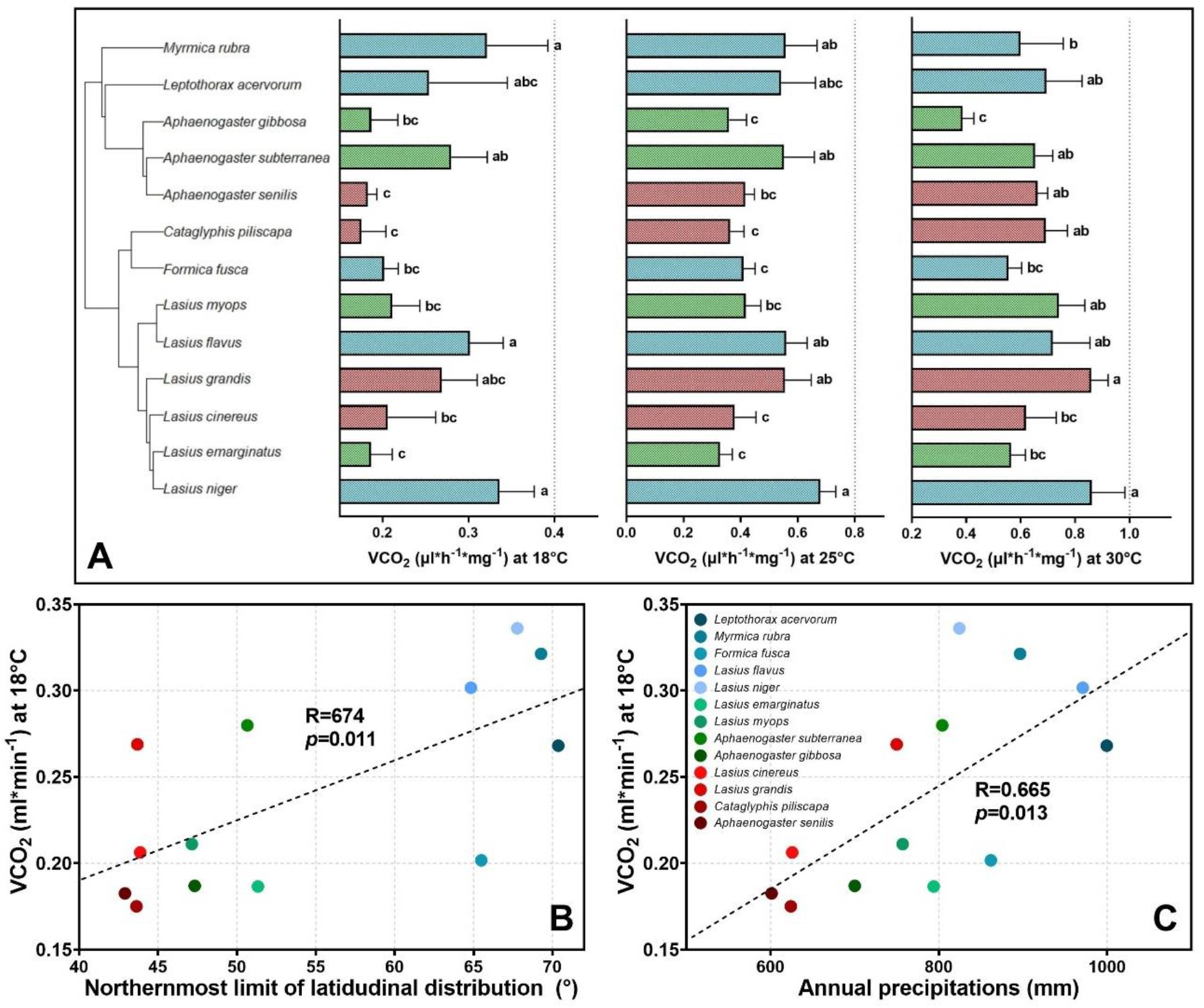
Standard metabolic rates at lower temperatures track species distribution. Species are color coded as being either Mediterranean (red) or extending their range towards intermediate (green) or northward (blue) latitudes within continental Europe. **A**. Averaged mass-corrected SMRs at 18, 25 and 30°C (different lowercase letters indicate significant differences between treatments; one-way ANOVA and Tukey *post hoc* test, *p*<0.05). **B**. Phylogenetically-corrected correlation between Northernmost limits of latitudinal distribution and mass-corrected SMRs at 18°C. Species extending their distribution northward displayed significantly higher respiration rates at 18°C (r=0.674, *p*=0.011). **C**. Phylogenetically-corrected correlation between annual precipitation and mass-corrected SMRs at 18°C (r=0.665, *p*=0.013).

**Table 2.** Pearson’s correlation coefficient (r) of the regressions between climatic variables averaged over species distribution, cold-tolerance, and respiration rates at three temperatures for the 13 species included in our analysis. The strength of these correlations is presented in shades of red (r values), and its statistical significance in shades of blue (*p* values). Overall, there was a very strong correlation between soil northernmost limits of latitudinal distributions (r=0.674, *p*=0.011, Fig.3B), annual precipitation (r=-0.665, *p*=0.013, Fig.4C), and respiration rates at colder, but not warmer, temperatures.

| Variables | VCO<sub>2</sub> (μl·h<sup>-1</sup>·mg<sup>-1</sup>) |  |  |  |  |  |
| --- | --- | --- | --- | --- | --- | --- |
|  | 18°C |  | 25°C |  | 30°C |  |
|  | r | p | r | p | r | p |
| Northernmost latitudinal distribution | 0.674 | 0.011 | 0.597 | 0.031 | 0.085 | 0.781 |
| Annual precipitation | 0.665 | 0.013 | 0.575 | 0.039 | 0.130 | 0.673 |
| Mean soil temperature diurnal range | -0.638 | 0.018 | -0.556 | 0.048 | -0.281 | 0.351 |
| Max soil temperature | -0.608 | 0.027 | -0.479 | 0.098 | -0.149 | 0.627 |
| Annual mean soil temperature | -0.588 | 0.034 | -0.432 | 0.140 | -0.043 | 0.89 |
| Min soil temperature | -0.360 | 0.226 | -0.193 | 0.527 | -0.185 | 0.544 |
| CT<sub>min</sub> | -0.204 | 0.503 | -0.103 | 0.738 | -0.296 | 0.325 |
| Southernmost latitudinal distribution | -0.127 | 0.678 | -0.012 | 0.968 | -0.089 | 0.773 |

### Lower Q_10-SMR_ at warmer temperatures for higher-latitude species

Q_10-SMR_ values were calculated for species between two intervals (18/25°C and 25/30°C; Fig. S.3). We then examined the putative correlations between Q_10-SMRs_ and absolute limits of latitudinal distribution, climatic variables averaged over species’ distributions, cold-tolerance, and respiration rates (Table 3). Overall, there were considerable differences in correlative strength between variables and the two temperature intervals of Q_10-SMR_. The best predictors of Q_10-SMR (25/30°C)_ were northernmost limits of latitudinal distribution (r=-0.678, *p*=0.011, Fig. S3B) as well as annual precipitation (r=-0.587, *p*=0.035, Fig. S3C). The best predictor of Q_10-SMR (18/25°C)_ was min soil temperature (r=0.530, *p*=0.062), although this was not statistically significant. Finally, we also observed a trend for a positive correlation between CT_min_ and Q_10-SMR (25/30°C)_ (r=0.521, *p*=0.068, Table 3). Overall, this indicates that species extending their range northward had significantly lower Q_10-SMR_ in the 25-30°C temperature interval, and displayed a shallower slope of the SMR TPC in that range, suggestive of a closer proximity with peak SMRs.

**Table 3.** Pearson’s correlation coefficient (r) of the regressions between climatic variables averaged over species distribution, limits of latitudinal distribution, cold-tolerance, respiration rates, and Q_10-SMR_ values for the 13 species included in our analysis. The strength of these correlations is presented in shades of red (r values), and its statistical significance in shades of blue (*p* values). Overall, there was a strong correlation between Q_10-SMR (25/30°C)_ and northernmost limits of latitudinal distribution, *p*=0.011, Fig. 3B) as well as annual precipitation (r=-0.587, *p*=0.035, Fig. S3C). There was no statistically significant association between climatic variables and Q_10-SMR (18/25°C)_.

| Variables | Q<sub>10-SMR</sub> |  |  |  |
| --- | --- | --- | --- | --- |
|  | 25/30°C |  | 18/25°C |  |
|  | r | p | r | p |
| Northernmost latitudinal distribution | -0.678 | 0.011 | -0.279 | 0.355 |
| Annual precipitation | -0.587 | 0.035 | -0.363 | 0.223 |
| CT<sub>min</sub> | 0.521 | 0.068 | 0.316 | 0.293 |
| Annual mean soil temperature | 0.505 | 0.078 | 0.375 | 0.206 |
| Max soil temperature | 0.415 | 0.158 | 0.411 | 0.162 |
| Min soil temperature | 0.390 | 0.188 | 0.530 | 0.062 |
| Mean soil temperature diurnal range | 0.310 | 0.302 | 0.260 | 0.391 |
| Southernmost latitudinal distribution | -0.042 | 0.895 | -0.338 | 0.258 |

### The association between cold tolerance and respiration rates at lower temperature predicts species’ distribution and climatic niche

Principal Component Analysis (PCA) of the variables included in our analysis highlighted that 55% of the variance found in our dataset could be explained alongside a single dimension to which most climatic variables contributed strongly, indicating that these were largely encompassed within a single “Northern climatic axis” (in other word, higher-latitude regions in Europe also display lower annual mean soil temperatures, lower min/max temperatures of the coldest/warmest months, and receive more precipitations, Fig. 4A). The two physiological variables best aligned with this axis were cold-tolerance (CT_min_,9.23%) and standard metabolic rates at 18°C (SMR_18°C_,8.12%, Fig. 4B). The two climatic variables best aligned with this axis were northernmost limits of latitudinal distribution (LatN, 13.08%) and annual mean soil-surface temperatures (Amt, 12.76%, Fig 4B). Following this, both LatN and Amt were further included in complementary model as representative climatic factors, to test the relative contribution of physiological variables (CT_min_, SMR_18°C_ and Q_10-SMR (25/30°C)_) to distribution and climatic niche. In line with the PCA results, the best predictive Phylogenetic Generalized Least Squares models (PGLS) for LatN and Amt performed included in each case CT_min_ and SMR_18°C_ as factors only (based on Akaike information criterion (AIC) scores; Table 4). Finally, we tested for a potential interactive effect between these two best physiological predictors (CT_min_ and SMR_18°C_) and found none (Table S.1). Details of PGLS models including CT_min_ and SMR_18°C_ can be found in table S.1.

**Table 4.** Akaike information criterion (AIC) weights for each Phylogenetic Generalized Least Squares model (PGLS) model tested to predict species’ northernmost limit of latitudinal distribution (LatN) and annual mean soil-surface temperatures averaged over their distribution range (Amt). In each case, inclusion of cold-tolerance (CT_min_) and respiration rates at 18°C (VCO_2 **(18**°C)_) in models only supported the best predictions (AIC score of 0.51 and 0.56, in green respectively).

| LatN | Variables included | Fit | Delta | AIC score |
| --- | --- | --- | --- | --- |
|  | CT<sub>min</sub> | 88.22 | 7.71 | 0.01 |
|  | SMR<sub>18°C</sub> | 94.62 | 14.10 | 0.43*10<sup>-4</sup> |
|  | Q<sub>10</sub>-SMR (25/30°C) | 95.70 | 15.18 | 0.2510<sup>-4</sup> |
|  | CT<sub>min</sub> + SMR<sub>18°C</sub> | 80.51 | 0.00 | 0.51 |
|  | CT<sub>min</sub> + Q<sub>10</sub>-SMR (25/30°C) | 84.30 | 3.78 | 0.07 |
|  | SMR<sub>18°C</sub> + Q<sub>10</sub>-SMR (25/30°C) | 95.61 | 15.09 | 0.26*10<sup>-4</sup> |
|  | CT<sub>min</sub> + SMR<sub>18°C</sub> + Q<sub>10</sub>-SMR (25/30°C) | 80.97 | 0.45 | 0.40 |
| Amt | Variables included | Fit | Delta | AIC score |
|  | CT<sub>min</sub> | 48.14 | 2.78 | 0.14 |
|  | SMR<sub>18°C</sub> | 61.96 | 16.60 | 0.01*10<sup>-3</sup> |
|  | Q<sub>10</sub>-SMR (25/30°C) | 60.73 | 15.36 | 0.02*10<sup>-2</sup> |
|  | CT<sub>min</sub> + SMR<sub>18°C</sub> | 45.36 | 0.00 | 0.56 |
|  | CT<sub>min</sub> + Q<sub>10</sub>-SMR (25/30°C) | 49.38 | 4.02 | 0.07 |
|  | SMR<sub>18°C</sub> + Q<sub>10</sub>-SMR (25/30°C) | 62.21 | 16.85 | 0.01*10<sup>-4</sup> |
|  | CT<sub>min</sub> + SMR<sub>18°C</sub> + Q<sub>10</sub>-SMR (25/30°C) | 47.18 | 1.82 | 0.22 |

**Fig. 4.**
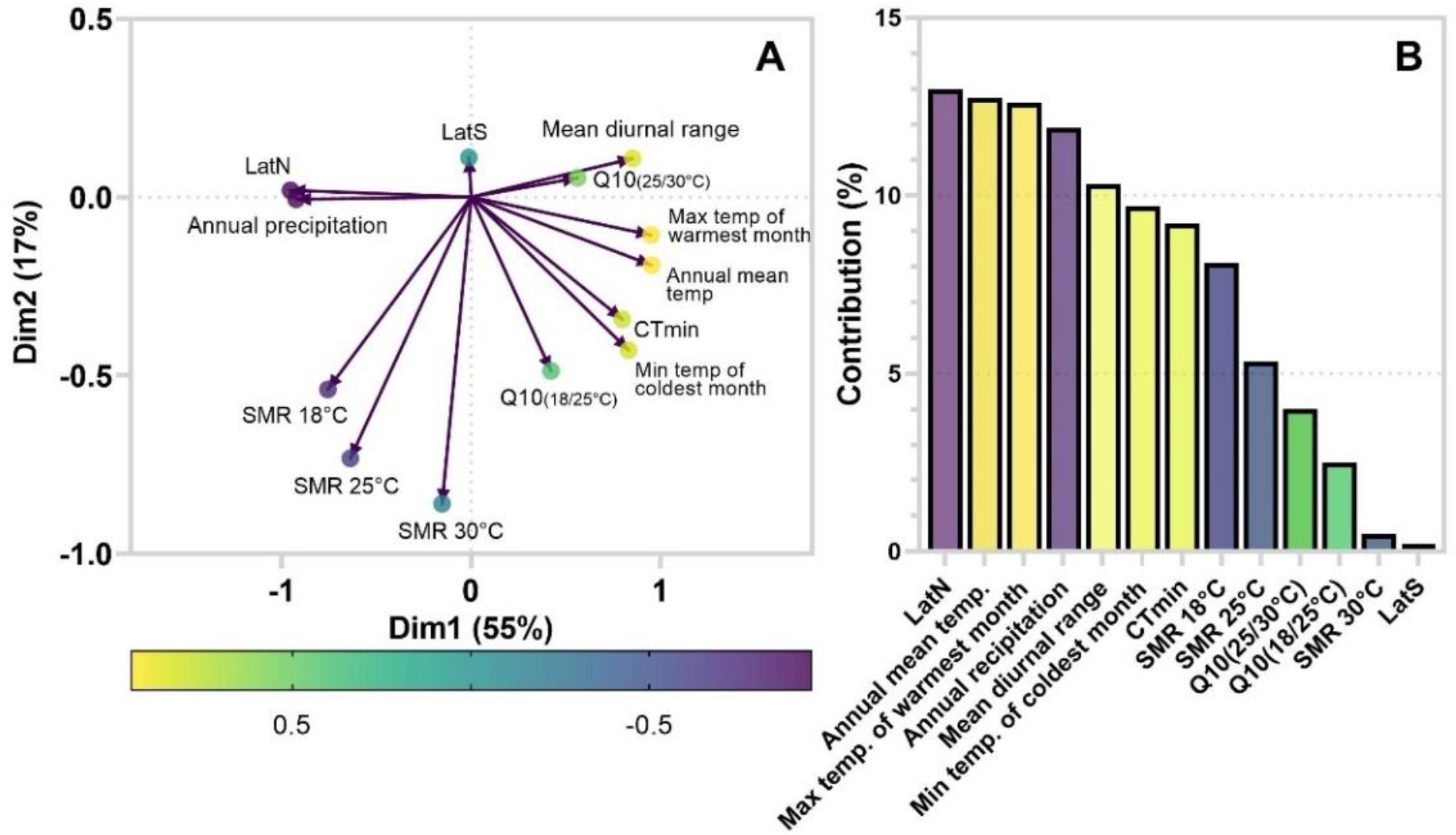
Principal Component Analysis of the physiological and climatic variables included in our dataset. **A**. A single dimension (Dim1) explained 55% of the variance found, in which most climatic variables contributed strongly, indicating that these were largely encompassed within a single “Northern climatic axis”. **B**. Variables ranked by their relative contribution to Dim1. Northernmost limits of latitudinal distribution (LatN) and annual mean temperatures (Amt) were the climatic variable respectively contributed the most to this dimension (13.08 and 12.76%, respectively). Color code ranges from strong positive (yellow) to strong negative (purple) association of variables with Dim1.

## Discussion

Understanding the key adaptations that enable ectotherms to thrive in cold climates have been a topic under scrutiny for decades [8, 11, 47]. In particular, cold-tolerance traits have been found as an unambiguous hallmark of colder-climate adaptation in ectotherms, that display strong associations with both altitudinal/latitudinal distribution as well as climatic niche in most species, including insects [17-21]. In contrast, the potential contribution of metabolic compensation as another evolutive tool for ectotherms to cope with colder temperatures has so far received ambiguous support [4, 5, 25]. Analytical and methodological biases when attempting broad inferences from multiple dataset have been among the fiercest critiques raised against the detection of MCA/MHH at larger scale in ectotherms [5]. Mixed results still emerge when considering studies individually and/or within phylogenetic groups, with for example in invertebrates, metabolic compensation effects detected in freshwater insects [48], in tsetse flies [7] or in woodlice [13], but only partial support for the MCA in snails [15], and an absence of support in *Drosophila* [16, 22, 49]. It is likely that such discrepancies capture some degree of biological truth: namely, that different invertebrates might have evolved different strategies to cope with high seasonality and low temperatures. Accordingly, the relevance of the MCA may be contingent on other life-history traits and therefore not unequivocally shared by all taxa.

In this study, data on SMRs at three thermal regimes in a methodologically and phylogenetically consistent framework provided strong support towards the occurrence of latitudinal metabolic compensation in ants. Species adapted to colder annual-mean soil temperatures and distributed at higher latitudes also displayed on average higher respiration rates at the lower (18°C, Fig. 3A) but not at the warmer (30°C) range of temperature tested. This pattern is highly consistent with the MCA hypothesis. Accordingly, SMRs at lower temperatures, but not warmer, were significant predictors of species latitudinal distribution and environmental soil temperatures (Table 2). Finally, we also report a negative relationship between northernmost latitude and Q_10-SMR (25/30°C)_ (Table 3, Fig. S3B), that is supportive of the MHH. Only a limited number of studies have probed for metabolic compensation effects specifically in the ant model. Although no links with cold-tolerance and distribution at broader scales were made, these also yielded a rather consistent support for metabolic compensation to colder environments [6, 50-53], for example by comparing populations of *Pogonomyrmex* [52], *Camponotus* [53], or *Aphaenogaster* [6] from different altitudes, or by comparing north-American species of ants from different climatic areas [51]. Here, leveraging the formal framework provided by our contemporary understanding of the MCA and MHH in ants [6], and using current biogeographical tools within a larger phylogenetical system, our data provides ongoing support for the occurrence of metabolic compensation specifically alongside a latitudinal axis in ants. Disentangling the relative contribution of plasticity at the population level versus potential interspecific adaptative shifts in this system cannot be addressed through our experimental design, which represent an inherent limitation of choosing to work with single populations of larger numbers of species. However, in light of our own and previous data, it should now be well accounted for that ants can indeed achieve metabolic compensation to colder climates either through phenotypic plasticity [6, 50, 52, 53], interspecific adaptative shifts [50-52] or, likely, through a combination of both.

The reduced metabolic thermal sensitivity observed in the 25-30°C range of temperatures for temperate species (Fig S.3, Table 3) could have ecological implications, considering the expected increases in mean temperatures and extreme climatic events under the current and future global warming [54]. The displacement of the SMR TPC towards cooler ranges of temperature through metabolic compensation in ectotherms might trade-off with metabolic performances at warmer ranges (Fig. 1) [26]. In corroboration with this, our data suggests that higher latitude species are indeed closer to peak SMRs than their Mediterranean relatives in the 25-30 °C range of body temperatures, which could indicate that these species might get further away of optimal temperatures for important traits such as growth and/or foraging. However, extrapolating to what degree metabolic compensation in colder-adapted species might trade-off with overall fitness at warmer temperatures, to support the documented effect of climate change on ant-communities [55-59] should be approached carefully. This is because operative temperatures (*i*.*e*. the steady state body temperature after accounting for all source of heat-transfer [60]) are not a passive function of climatic temperatures in ants, but are heavily regulated through foraging and nesting behaviors [61, 62]. In addition, species often occupy distinct thermal niches in their respective environments [63]. As a result, a warmer climate will not impact all ants species equally. For example, it is often observed that ants with cooler optima for traits such as foraging tend to simply behaviorally avoid too warm conditions [64-68]. Our data nonetheless suggests that a substantial part of temperate ant species may reach a tipping point in the 25-30°C range of body temperatures, where the performance of some critical life-history traits might start declining. It is thus tempting to hypothesize that a warming climate will open more windows of opportunity for species physiologically better equipped to deal with warmer temperatures in temperate regions, thus ultimately outcompete their cold-adapted relatives, pushing them towards cooler niches [55-59].

Finally, and considering the apparent importance of metabolic compensation to adapt to cold climates in our comparative analysis of ant species, we aimed at dissecting its relative contribution to associate specific traits to species distribution. As discussed above, cold-tolerance is typically a strong correlate of species distribution in insects, including ants [17-21], which remains well supported by our own analysis (Table 1, Fig. 2). However, we also show that CT_min_ and SMR_18°C_ align considerably on the same dimension as most climatic factors (Fig. 4A), and that the combination of both cold-tolerance and MRs at lower temperatures supports better prediction regarding species biogeography than either one alone (Table 4). This highlights the importance of accounting for metabolic rates when inferring species’ climatic niches in ants. The absence of a direct correlation between MRs and CT_min_ also offers interesting perspectives. It suggests that either these two traits operate independently, or that the range of temperatures where this relationship might become apparent is located well below the 18°C threshold that is roughly associated with the lower permissive temperature for foraging activity of similar models [67, 68]. One could also hypothesize that MRs and CT_min_ might represent to different, complementary aspects of cold-climate adaptation. Increased cold-tolerance could for example help surviving temperature extremes, while metabolic compensation could enable resource acquisitions during longer term, but suboptimal, conditions.

In conclusion, we report here a clear metabolic compensation effects within 13 ant species distributed across a wide latitudinal gradient in western Europe, highly supportive of the Metabolic Cold Adaptation and Metabolic Homeostasis Hypothesis. We further show that the cold-tolerance correlate very well with species distribution, but also that the combination of cold-tolerance and metabolic rates at lower temperatures is even better at predicting species limit of distribution range and climatic niche than either one alone. Overall, these results highlight that evolution to colder-climate is a complex trait in ants, which, potentially at the expense of metabolic performance at warmer temperatures, has leveraged shifts in both cold-tolerance as well as metabolic rates to allow them to thrive under northern latitudes.

## Acknowledgements

We thank L. Jørgensen for her help in taking care of captive ant colonies, and C. Damsgaard for his help with phylogenetic analysis. We are further grateful to K. Becq, L. Vanbegin and S. Vanbegin for their help with field sampling, as well as to X. Espalader for his help with *Lasius*-niger group species identification.

## Authors’ contributions

Q.W. and J.O. conceived and planned the study. Q.W. collected samples. Q.W. performed lab experiments, and Q.W. M.Ø. and H.M. analysed the data. All authors contributed to drafting the article, approved the final published version, and agree to be held accountable for all aspects of the work.

## Data availability

All data are included in the manuscript and Supporting Information.

