## Supplementary figures and tables for "Cold Comfort: metabolic rate and tolerance to low temperatures predict latitudinal distribution in ants"

**Electronic supplementary material**

[a]Department of Biology, Aarhus University, 8000 Aarhus C, Denmark

[b]Department of Chemistry and Bioscience, Aalborg University, 9220 Aalborg E, Denmark

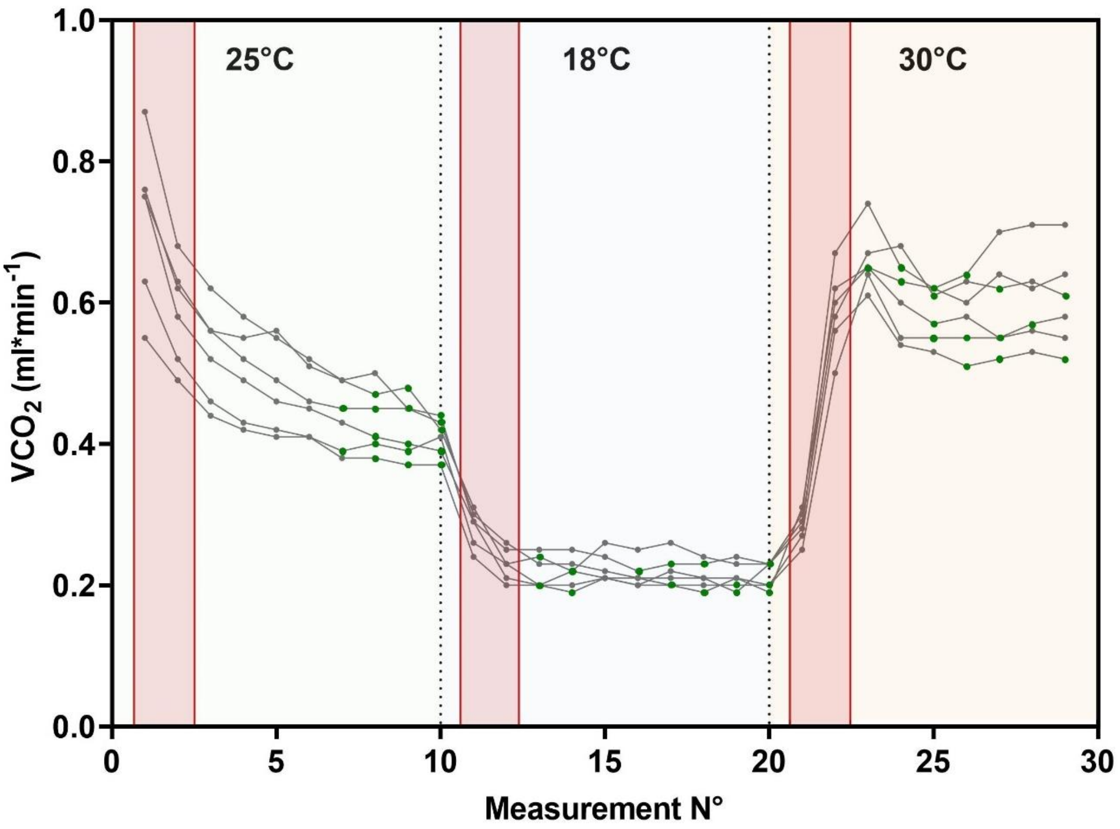


**Figure S1.** Representative respirometry run (*Formica fusca*) recording non-mass corrected respiration rates of workers at 3 temperatures over a 24h period. Each curve represents a chamber (*i.e.,* replicate) containing 20 workers. Each measurement was taken at a 48min interval (*i.e.,* $CO_2$ was left to accumulate solely from workers´ respiration for 48min before each new measurement). The temperature inside the setup was programmed to shift from 25 to 18 and then 30°C at 8h intervals (ca. 480 min, 10 measurements). As it took approx. 90min for the setup to equilibrate between temperature shifts, the first two measurements at each temperature (red highlights, ca. 90min) were therefore systematically excluded from the analysis to allow workers to settle after each transition. Finally, only the three lowest values for each replicate (green dots) were averaged at each trial temperature and taken into account for subsequent analysis.

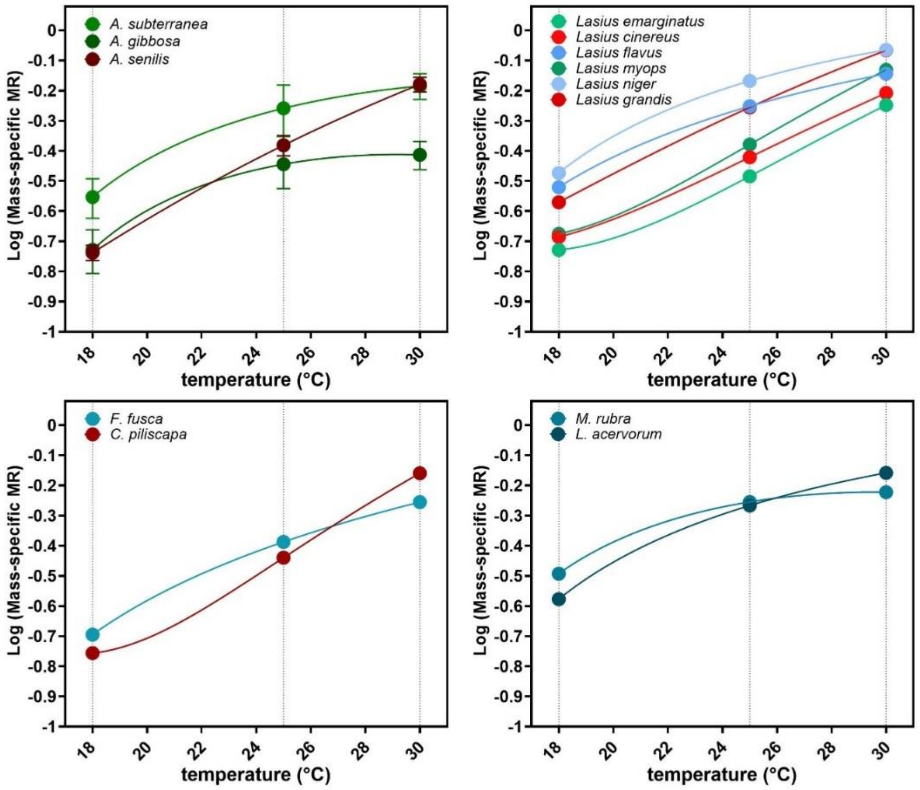


**Fig. S.2.** Metabolic rate curves for the 13 species included in our analysis, split into 4 groups for better visualization. Respiration rates were assayed at three temperatures (18, 25 and 30°C), and fitted with quadratic functions following Shik *et al.*, 2019 [1] (please see the latter reference for the use of quadratic rather than linear functions to fit thermal performance curve of the SMR in ants). Groups are further color coded as either representing curves for Mediterranean species (shades of red), or species extending their range to either intermediate (shades of green) or northward (shades of blue) latitudes within continental Europe. **A.** Metabolic rate curves for *Aphaenogaster* species. **B.** Metabolic rate curves for *Lasius* species. **C.** Metabolic rate curves for species within the Formicini tribe. **D.** Metabolic rate curves for *Leptothorax acervorum* and *Myrmica rubra*. Overall, species extending their range northward (blue) exhibited higher respiration rates at 18 and 25, but not 30°C. On average, the slope of the quadratic function between points ($Q_{10\text{-SMR}}$) decreased between 25 and 30°C for northern species (blue) as compared to Mediterranean ones (red). The stop flow respirometry system as used in the present study was demonstrated in *Drosophila* to produces reliable SMR estimates that are generally slightly lower than measurements obtained using more common open flow protocols[2, 3]. Consistently, our mass-specific metabolic rates and $Q_{10\text{-SMR}}$ values appear on the lower end of what was previously documented for ants [1, 4-6].

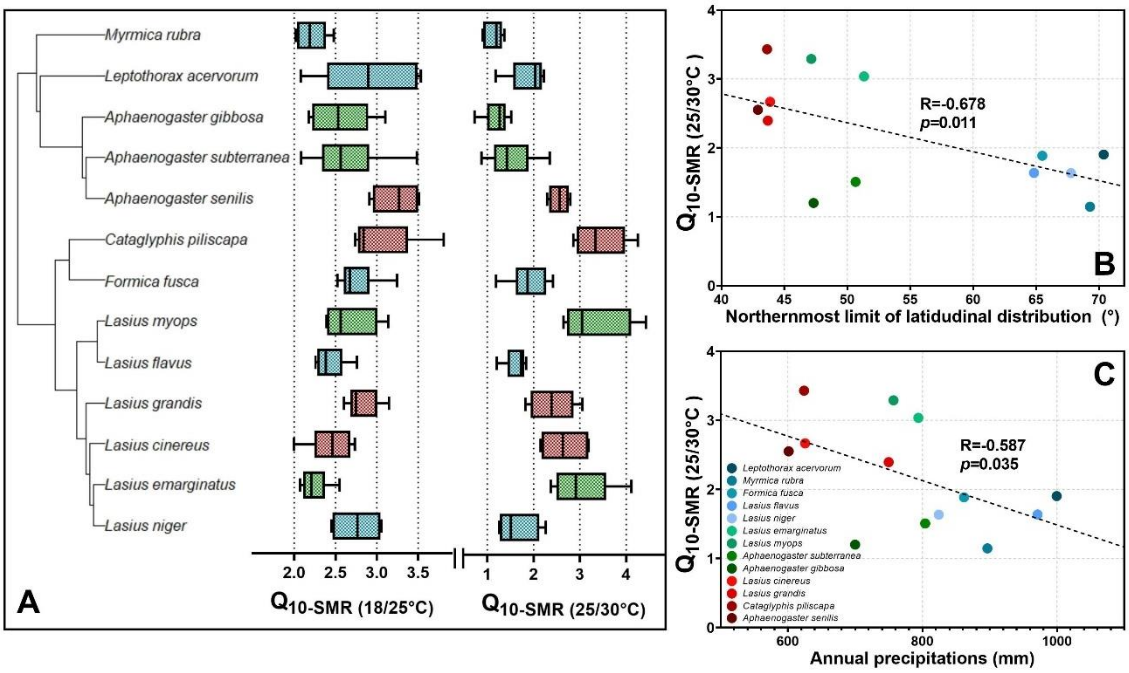


**Figure S.3.** $Q_{10\text{-}SMR}$ values Averaged over 3-6 respirometry chambers per species for each temperature interval tested. $Q_{10\text{-}SMR}$ values decreased at warmer intervals for cold-adapted species. **A.** $Q_{10\text{-}SMR}$ values for the 13 species included in our analysis calculated for the 18/25 and 25/30°C intervals. **B.** Correlation between $Q_{10\text{-}SMR\ (25/30°C)}$ and species' northernmost limit of latitudinal distribution. Species distributed northward displayed reduced $Q_{10\text{-}SMR}$ at 25/30°C (r=-0.678, *p*=0.011 also see Fig. S.2). **C.** Phylogenetically-corrected correlation between annual precipitations and $Q_{10\text{-}SMR\ (25/30°C)}$ (r=-0.587, *p*=0.035).

| | | $CT_{min}$ | | $VCO_{2(18°C)}$ | | $CT_{min}$ x $VCO_{2(18°C)}$ | |
|---|---|---|---|---|---|---|---|
| **Model N°** | **Model** | ***t* value** | ***p* value** | ***t* value** | ***p* value** | ***t* value** | ***p* value** |
| 1 | LatN ~ $CT_{min}$ | -4.689 | >0.001* | / | / | / | / |
| 2 | LatN ~ $SMR_{(18°C)}$ | / | / | 3.0281 | 0.011* | / | / |
| 3 | LatN ~ $CT_{min}$ + $SMR_{(18°C)}$ | -4.951 | >0.001* | 3.332 | 0.007* | / | / |
| 4 | LatN ~ $CT_{min}$ + $SMR_{(18°C)}$+ $CT_{min}$* $SMR_{(18°C)}$ | -1.750 | 0.114 | 2.987 | 0.015* | 0.990 | 0.349 |
| 5 | Amt ~ $CT_{min}$ | 5.932 | >0.001* | / | / | / | / |
| 6 | Amt ~ $SMR_{(18°C)}$ | / | / | -2.413 | 0.034* | / | / |
| 7 | Amt ~ $CT_{min}$ + $SMR_{(18°C)}$ | 5.657 | >0.001* | -2.108 | 0.061 | / | / |
| 8 | Amt ~ $CT_{min}$ + $SMR_{(18°C)}$ + $CT_{min}$* $SMR_{(18°C)}$ | 2.714 | 0.023* | -2.870 | 0.018* | -1.76 | 0.111 |

**Table S.1**. Phylogenetic Generalized Least Squares model (PGLS) models predicting species' northernmost limit of latitudinal distribution (LatN) and annual mean soil-surface temperatures averaged over their distribution range (Amt), and including cold-tolerance ($CT_{min}$), standard metabolic rates at 18°C ($SMR_{(18°C)}$) and the interaction between the two as factors. We detected no significant interactions between cold tolerance and respiration rates at 18°C (models 4, 8). Models 3 and 7 overall supported the best prediction of LatN and Amt (Table 4).
